# Comparative analysis of genome-wide DNA methylation identifies patterns that associate with conserved transcriptional programs in osteosarcoma

**DOI:** 10.1101/2020.04.29.068155

**Authors:** Lauren J. Mills, Milcah C. Scott, Pankti Shah, Anne R. Cunanan, Archana Deshpande, Benjamin Auch, Bridget Curtin, Kenneth B. Beckman, Logan G. Spector, Aaron L. Sarver, Subbaya Subramanian, Todd A. Richmond, Jaime F. Modiano

## Abstract

Osteosarcoma is an aggressive tumor of the bone that primarily affects young adults and adolescents. Osteosarcoma is characterized by genomic chaos and heterogeneity. While inactivation of tumor suppressor p53 *TP53* is nearly universal other high frequency mutations or structural variations have not been identified. Despite this genomic heterogeneity, key conserved transcriptional programs associated with survival have been identified across human, canine and induced murine osteosarcoma. The epigenomic landscape, including DNA methylation, plays a key role in establishing transcriptional programs in all cell types. The role of epigenetic dysregulation has been studied in a variety of cancers but has yet to be explored at scale in osteosarcoma. Here we examined genome-wide DNA methylation patterns in 24 human and 44 canine osteosarcoma samples identifying groups of highly correlated DNA methylation marks in human and canine osteosarcoma samples. We also link specific DNA methylation patterns to key transcriptional programs in both human and canine osteosarcoma. Building on previous work, we built a DNA methylation-based measure for the presence and abundance of various immune cell types in osteosarcoma. Finally, we determined that the underlying state of the tumor, and not changes in cell composition, were the main driver of differences in DNA methylation across the human and canine samples.

**Significance:** This is the first large scale study of DNA methylation in osteosarcoma and lays the ground work for the exploration of DNA methylation programs that help establish conserved transcriptional programs in the context of different genomic landscapes.

## Introduction

Osteosarcoma is a rare disease, there are fewer than 1,000 cases diagnosed in the U.S. each year, mostly in children and adolescents^1^. However, these numbers fail to convey the impact that the disease has on patients, their families, caregivers, and the extended community due to its significant morbidity and years of life lost. Recent work is starting to increase our fundamental understanding of osteosarcoma^2^ but more than half of patients still relapse and die from metastatic disease within 10 years^1,3^. Osteosarcoma has been reported in every vertebrate species^4^. It is as rare in most animals as it is in humans, and when it occurs, it is most common in the axial skeleton^4^. Dogs are a notable exception. Osteosarcoma is extremely common in large and giant dogs, and similar to children the disease occurs most frequently in the appendicular skeleton^4,5^

Many studies have evaluated the genomic landscape of osteosarcoma in humans and in animal models. The heterogeneity of this disease is remarkable, both within and among species^5–10^. Loss of function of the *TP53* gene seems to be a nearly universal event in spontaneous osteosarcoma, thus it might be causally related to the chaotic genomes that are characteristic of this condition^7^. Aside from *TP53* loss of function mutations, recurrent genomic aberrations are rare within species and even more infrequent between species. But unlike its highly heterogeneous mutational landscape, the transcriptional programs that characterize human and canine osteosarcoma are highly conserved^8,11^. More specifically, one key transcriptional program defined subsets with higher (or lower) rates of tumor cell proliferation and turnover, inferred from the expression of gene clusters associated with cell cycle progression, mitosis, DNA damage repair, and chromosomal instability^8,11,12^. We created a method to quantify this expression using a gene cluster expression summary score (GCESS)^8^ and showed that this GCESS was inversely associated with overall survival in both dogs and humans. The other salient conserved transcriptional programs defined subsets associated with abundance of immune and inflammatory cells in the microenvironment, inferred from the expression of genes uniquely or predominantly expressed by cells of the innate and adaptive immune system^8^. Curiously, the immune GCESS were only predictive of survival time and metastasis in human patients and not in dogs.

The discrepancy between the highly heterogeneous mutational landscape of human and canine osteosarcomas with diverse putative genetic drives and their relatively homogeneous transcriptional landscape indicates these two sets of events are probably unrelated in both species. In other words, these tumors do not seem to be caused by driver events that activate or repress specific transcription. Instead, it is most likely that osteosarcomas in humans and in dogs are convergent entities where epigenetic controls of gene expression driven by selection ultimately give rise to the limited molecular pathways that achieve the tissue organization required to form osteosarcoma tumors. One such epigenetic control is DNA methylation, which in turn is a major determinant of chromatin accessibility.

For this study, we sought to determine the role of DNA methylation in establishing the conserved transcriptional programs observed in human and canine osteosarcoma. Our data show that, indeed, there are conserved modules of methylation in human and canine osteosarcomas. In spite of the apparent species differences, such conserved processes must reflect pathogenetically significant events that contribute to risk, progression, and therapeutic failure, and understanding their mechanisms will aid the development of better methods to identify risk and prognosticate progression, and ultimately more effective strategies for treatment, control, and eventually prevention.

## Results

### Genome wide DNA methylation patterns in human and canine osteosarcoma

Our objective was to establish mechanisms that control conserved transcriptional programs in human and canine osteosarcoma tissues. Targeted bisulfite sequencing was used to measure genome wide DNA methylation levels in 24 samples of human osteosarcoma and 44 samples of canine osteosarcoma. In order to focus the comparisons on conserved mechanisms that underlie the cellular and molecular organization of osteosarcoma, we only considered methylation measurements from homologous regions of the two genomes (54K human and 58K canine genomic regions, 19% and 33% of total measured regions respectively, Figure 1A). Weighted gene correlation network analysis (WGCNA)^13,14^ clustering was applied to the DNA methylation data to reduce complexity by identifying genomic regions with highly correlated DNA methylation profiles across the osteosarcoma samples in both species. WGCNA generates large clusters of genomic regions with highly correlated DNA methylation measurements across samples and produces a summary value for each cluster. Clusters were constructed using the same parameters for the human and canine samples and the same correlation cut off value was used to establish cluster membership. We thus condensed 12,165 unique methylation measurements from the human osteosarcoma samples into 43 clusters containing between 16 and 2,529 individual methylation measurements (Figure 1A). Five modules contained less than 100 measurements and 6 modules contained more than 1,000. Similarly, we condensed 6,099 unique methylation measurements from the canine osteosarcoma samples into 12 clusters containing between 139 and 2,532 individual methylation measurements (Figure 1A). The representative methylation value for each cluster, equivalent to the first principal component eigenvalue, is given in the heatmaps in Figure 1B and 1C. For both the human and canine clusters, methylation measurements from multiple chromosomes are clustered together indicating larger methylation programs that extend beyond methylation measurements in local genomic regions. While only methylation measurements from homologous genomic regions were used for clustering, the human and canine samples were clustered independently because we were interested in identifying methylation clusters that contain homologous genomic regions in the two species. The highest levels of region overlap were seen between human cluster ME1 and canine clusters ME1 (31.75%) and ME10 (33.26%), between human cluster ME5 and canine cluster ME3 (33.33%), and between human cluster ME7 and canine cluster ME8 (33.61%) (**supp table**). K-means clustering using the Euclidean distance of the summarized methylation values for each cluster was used to identify clusters that showed similar patterns across either the human or canine osteosarcoma samples, as indicated by the colored toe bars at the bottom of Figures 1B and 1C. The color scheme in Figures 1B and 1C is maintained throughout the rest of the manuscript and used as a reference for clusters. Some of the clusters share genomic regions across both species, as indicated by shared colors in the toe bars (blue, green, orange), while other clusters are species specific (red, light orange, yellow, purple, and black, Figures 1B, 1C, and 1D). We assigned the nearest gene (≤1,000 bp) to each methylation measurement and performed pathway analysis using the Reactome database for each methylation cluster. Not all methylation clusters resulted in significant pathway enrichment, but with the exception of the black group, all others had 1 or more individual groups of methylation marks with enriched functional pathways (Figure 1D and **supp table**). Predictably^13,14^, genes in the Reactome pathways “Transcriptional Regulation by TP53: (R-HSA-3700989), along with other pathways associated with signal transduction, were enriched in modules in the blue group, and genes in the “Transcriptional regulation by RUNX2 ‘’ and “Transcriptional regulation of pluripotent stem cells” pathways were enriched in modules in the green group in samples from both species.

**Figure 1:**
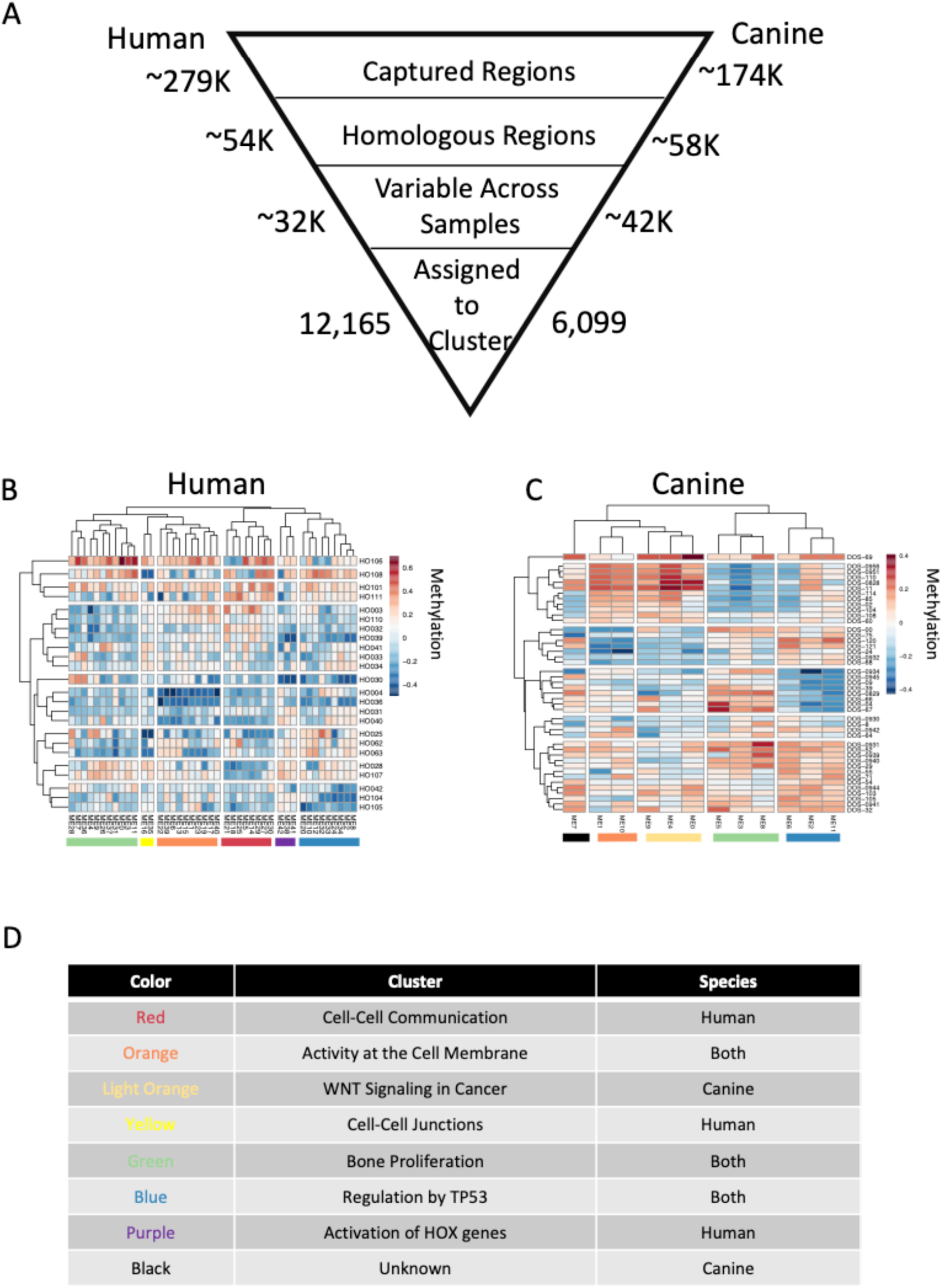
Genome wide DNA methylation patterns in canine and human osteosarcoma samples. Summary of analysis and the number of genomic regions included in each step (A). DNA methylation values were calculated at homologous genomic regions in canine (B, n=44) and human (C, n=24) primary osteosarcoma tumor samples. Genomic regions were clustered based on correlation in each species resulting in 12 canine clusters and 43 human clusters containing genomic regions that have highly correlated methylation values across all osteosarcoma samples. Heatmaps represent the weighted average DNA methylation value for the cluster in each sample. Red is a high methylation value and blue is a lower methylation value. Pathway analysis was performed using Reactome on the gene nearest (<=1,000bp) the underlying genomic region in each DNA methylation cluster. Clusters that share many of the same pathways are color coded the same in B and C and a summary of the pathways is given for each colored block in D. The unknown cluster did not have any significantly enriched pathways.

### Association between methylation clusters and osteosarcoma transcriptional programs

Previously, we identified shared transcriptional programs in human and canine osteosarcoma tissues that were associated with proliferation (cell cycle), presumably of the tumor cells that were inversely associated with overall survival in both species^8^. We also showed that the presence of immune cells (immune1 and immune2) in the microenvironment was directly associated with overall survival and time to metastasis, but only in human patients^8^. A subset of the samples used to identify those transcriptional programs were included in this study (human n = 16, canine n = 9). We calculated correlations between DNA methylation clusters and the summarized gene expression values for the expression groups from Scott et. al (2018) using the summarized methylation values for each cluster identified with WGCNA (Figure 2). In addition, we correlated DNA methylation to the age at diagnosis and overall survival time for the canine samples (Figure 2B). Figure 2 shows that the methylation clusters not only behave similarly when looking at the global DNA methylation patterns, but also when they are compared to downstream transcriptional programs. Specifically, the methylation clusters in the green group are positively correlated with cell cycle expression and inversely correlated with immune expression in both the dog and the human samples. In contrast the methylation clusters in the orange group, in both species, and the red group, in the human samples, show the opposite behavior and are inversely correlated with expression of cell cycle-associated genes and positively correlated with expression of immune-related genes.

**Figure 2:**
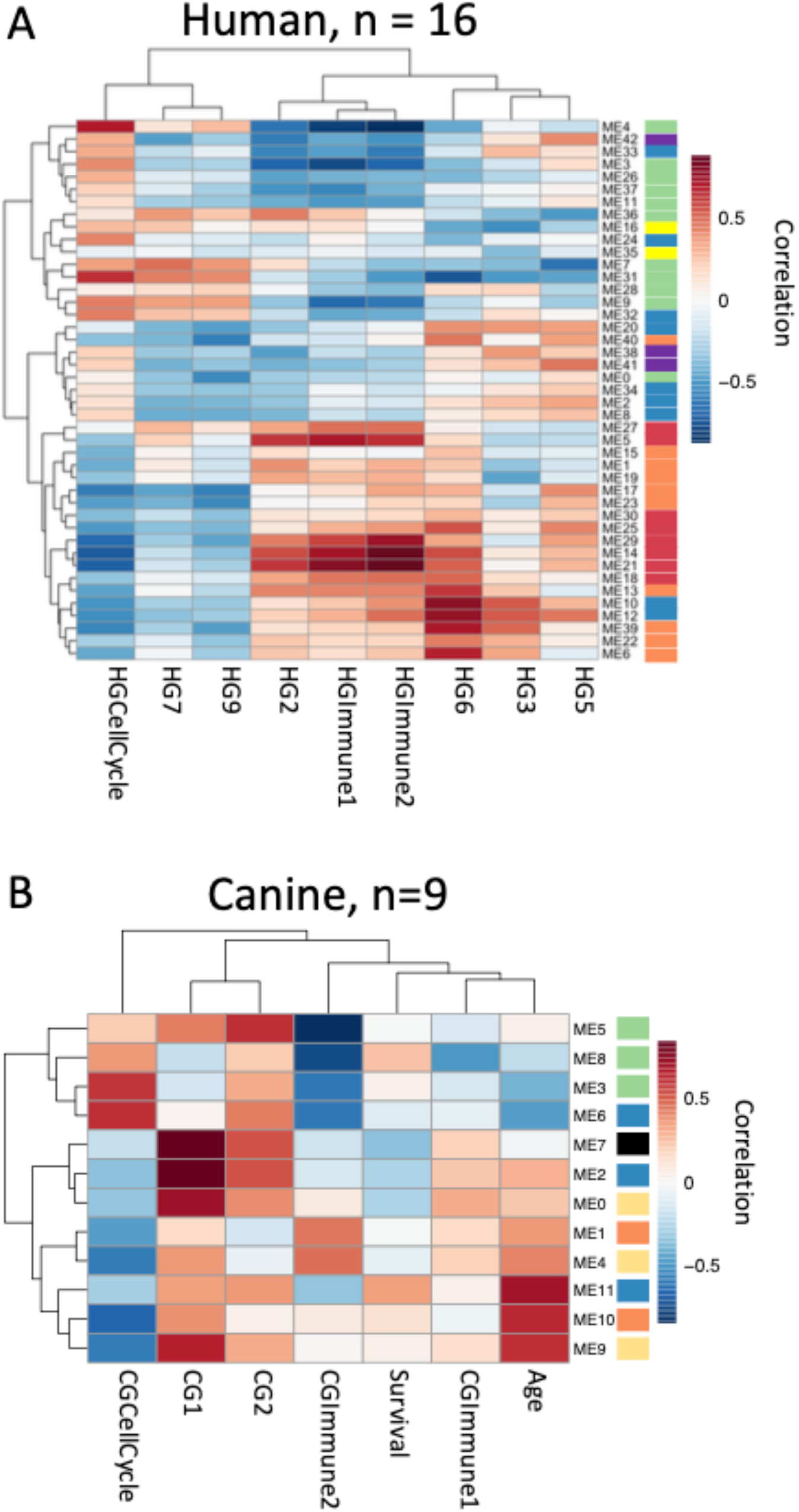
Correlation between methylation clusters and important expression programs in osteosarcoma. Correlations between methylation summary values for each DNA methylation cluster and previously described expression programs were calculated for canine (A, n=9) and human (B, n=16) samples where methylation and expression data was available. Correlations were also calculated for additional phenotypic data for canine samples. Heatmaps indicate Pearson correlation values between each DNA methylation cluster and phenotype where red is strong positive correlation and blue is strong negative correlation. Color bars next to module names indicate pathway analysis results and are the same as in the previous figure. HG = human GCESS, CG = canine GCESS from Scott et al 2018^8^.

### Immune cell abundance across osteosarcoma samples

The presence or absence of specific immune cell infiltrates is an important predictor of survival in many solid tumors^15^ including human osteosarcoma^8^. DNA methylation data, from 450K arrays, has been shown to be more effective at predicting the abundance of specific cell types in gold standard mixtures than expression data from RNA-seq^16^. Building on these previous methods, we obtained whole genome bisulfite sequencing data for 12 pure human cell populations from the BLUEPRINT project^17^. Whole genome bisulfite sequencing (WGBS) data were summarized to the same human/canine homologous regions used as input for WGCNA analysis. Cell populations of interest included10 immune cell populations as well as osteoclasts and mesenchymal stromal cells (MSC), which were used as surrogates for tumor cells as WGBS data for osteoblasts is not available in the public domain. We performed differential methylation tests between all pairs of pure cell populations, and the top 100 genomic regions with the largest differences in methylation levels were selected from each test and combined (1,312 unique regions) to generate a custom signature file in CIBERSORT (SFigure 1). Once generated, this custom signature file accurately predicted the cell type in computationally generated mixtures based on methylation levels (SFigure 2). WGBS data from known mixtures of canine cells were not available to further test predictions from this method, so we relied on the approach used to identify homologous genomic regions to convert canine methylation measurements to their human genome equivalents. We then applied the same signature file to both species and, as expected, MSCs had the highest signal across all samples (Figure 3A & 3B). The data also suggests that all other stromal and immune cell types that we interrogated were infrequent (low abundance) in human osteosarcoma samples (Figure 3A), while suggesting that monocytes, CD8+ T-cells, and osteoclasts were relatively more abundant in canine osteosarcoma samples (Figure 3B).

**Figure 3:**
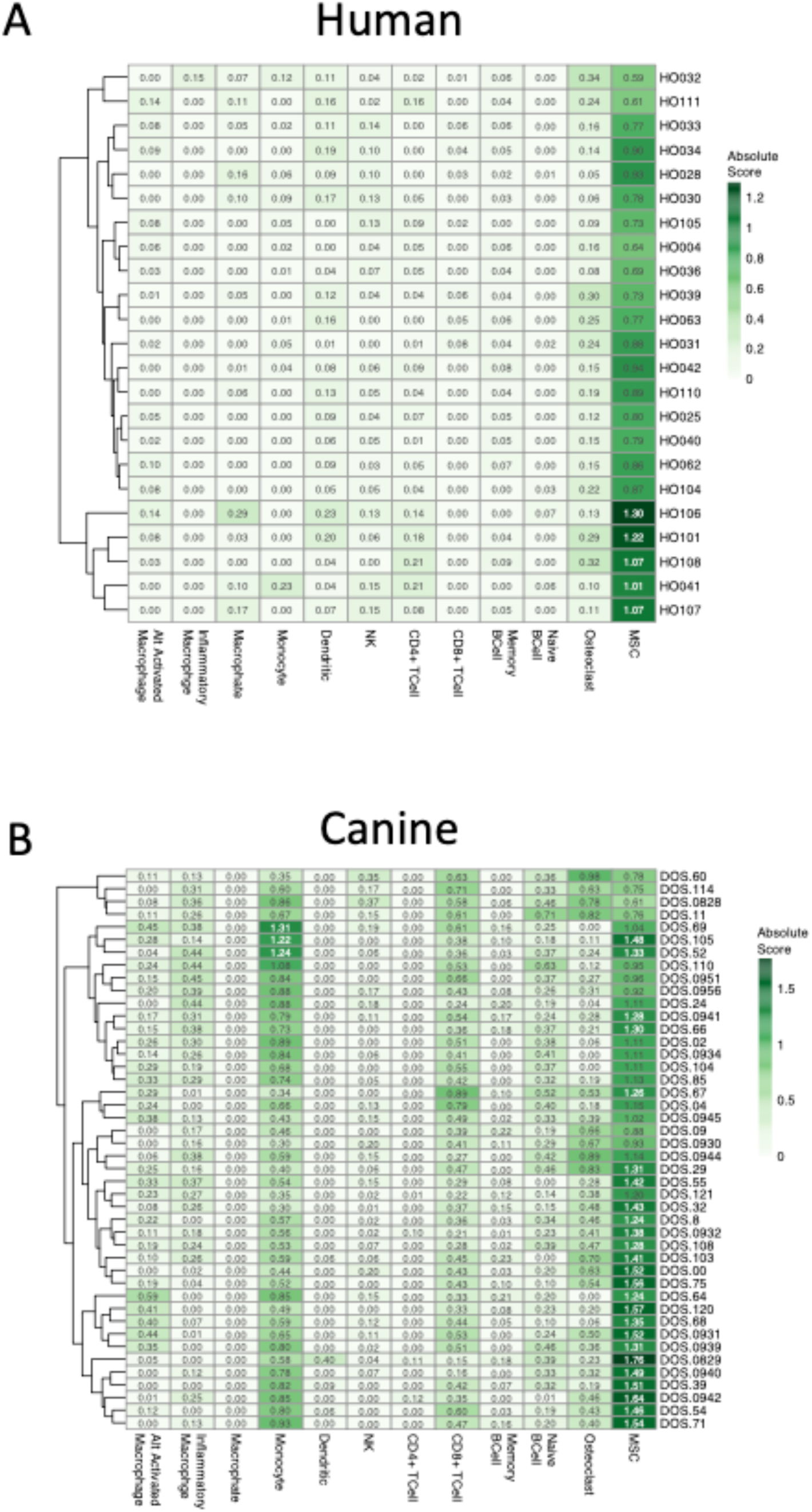
Immune cell abundance across osteosarcoma samples. WGBS data from pure human immune cell populations and mesenchymal stromal cells were obtained from the BLUEPRINT project. DNA methylation values were summarized to match those measured in our osteosarcoma samples and used to build a model to distinguish between each cell type. This model was used with CIBERSORT to measure the absolute cell abundances for each cell type in all of our osteosarcoma samples. Heatmap color and value in each cell indicate the abundance of each cell type in each sample for canine (A) and human (B) osteosarcoma tumors. Absolute mode was used to generate cell abundance scores and are not relative so columns will not sum to 1 (100%).

### Associations between global DNA methylation modules and predicted immune cell abundance

Global DNA methylation patterns are highly specific to cell type, though obtained through separate analysis, it is possible that the DNA methylation clusters identified via WGCNA were a result of the differences in the abundance of different cell types in the osteosarcoma samples. Though generated from the same starting regions (Figure 1A), there was a small overlap between the regions included in the WGCNA methylation clusters and the custom signature file used with CIBERSORT (352/12,165 from the human data and 147/6,099 from the canine data). To understand if any of our discovered methylation clusters were indirectly measuring immune cell abundance or the state of the tumor tissue, we calculated correlations between the summarized methylation values for each methylation cluster and predicted immune cell abundance scores (Figure 4). In both species, similar to the relationships between the methylation scores and transcription measures (Figure 2), methylation clusters of the same color tend to have similar relationships to the cell abundance measures (Figure 4). In the human data most of the methylation clusters showed similar, positive, correlations to MSC while they displayed a more varied behavior overall (Figure 1A) and against key transcriptional patters (Figure 2A). Interestingly, the orange and green clusters both have positive correlations to MSC abundance while they have opposite correlations, from each other, with the immune and cell cycle expression measures (Figure 2). In the canine data strong positive and negative correlations are seen between MSC, CD8+ T-cells and inflammatory macrophages. Also, in the canine data the opposite correlations are seen between the green and orange/light orange groups mirroring what is seen when compared to the transcriptional patters (Figure 2B). In the human data, immune cell abundance is very low and the strongest correlations are seen between the methylation modules and the MSC abundance measures. While a few of the methylation clusters may reflect the abundance of immune cells, many more seem to reflect the underlying states of the tumor itself. In the canine data, immune cell abundance is higher and more varied, and the same modules show strong correlations to both transcriptional measures of immune cell infiltration (Figure 2B) and methylation-based measures (Figure 4B), so more of the original methylation clusters (green clusters) may be reflective of cell abundance. But, many clusters do not show a strong correlation to any cell abundance measure and most likely reflect other aspects of the underlying tumor state.

**Figure 4:**
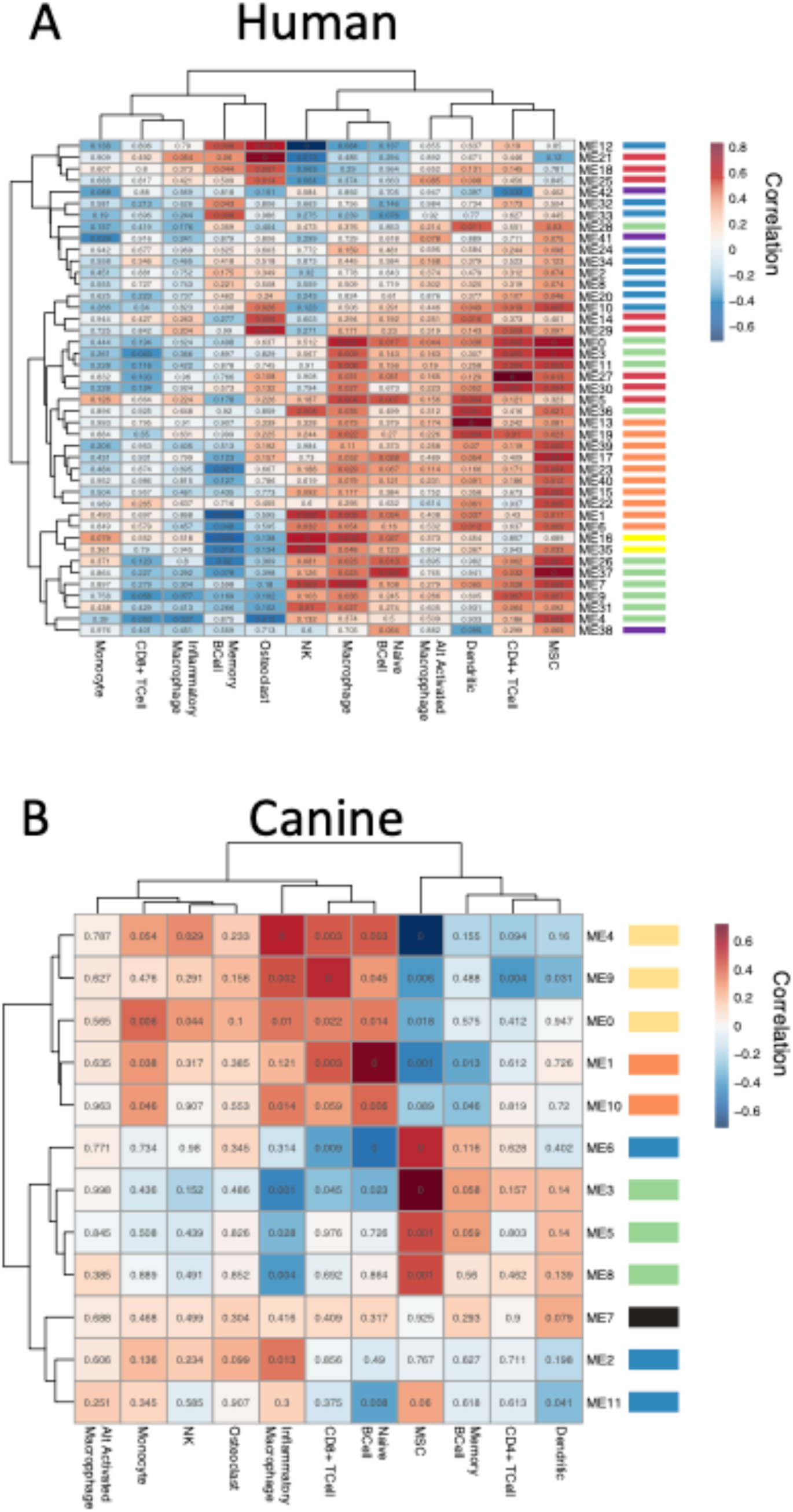
Correlation between methylation clusters and cell type abundances in OS. Correlations between methylation summary values for each DNA methylation cluster and CIBERSORT absolute cell type abundances were calculated for canine (A, n = 44) and human (B, n = 24) samples. Heatmaps indicate Pearson correlation values between DNA methylation cluster and cell type abundance where red is a strong positive correlation and blue is a strong negative correlation. Values in each cell are the Student asymptotic p-value of the given correlation based on the number of samples. Color bars next to module names indicate pathway analysis results and are the same as in the previous figures.

## Discussion

The molecular basis of osteosarcoma has received considerable attention during the last decade^18–26^. Several genetic mutations including p53, RB, cMYC, and RUNX2 have been strongly implicated in the development of osteosarcoma^7,24,27–38^ but our understanding of osteosarcoma epigenome is still limited^8–43^. The clinical outcomes continue to be dismal and have landed this disease among the “most wanted” for development of new, effective therapies. Elements of our failure to understand osteosarcoma pathobiology and treatment include tumor heterogeneity, a lack of robust prognostic factors, and the fact that current therapies ultimately fail to prevent relapse and/or metastasis for most patients. These challenges are also compounded by the *orphan disease* status of osteosarcoma.

Recently, we showed that transcriptional programs in human and canine osteosarcoma defined the proliferation and immune signature in the tumor microenvironment that are associated with aggressiveness and outcomes in osteosarcoma. Notably, our studies and other genomic studies have failed to identify recurrent translocation or mutational profiles associated with the transcriptional programs in human and canine osteosarcoma. In the absence of recurrent mutations the aberrant expression of pro-survival and metastatic genes noticed in osteosarcoma may be in part due to deregulation of microRNAs^39,40^. In addition to microRNAs the transcriptional programs may be regulated by epigenetic alterations^41–43^. Using genome-wide methylation profiles of human and canine osteosarcoma, here we show that transcriptional programs associated with cell proliferation and immune signature are associated with methylation patterns in both the species. Specifically, DNA methylation across the blue, green and orange clusters encompassing 1,773 genomic locations behave similarly between human and canine while other regions are species specific. Correlative analysis of genome-wide methylation patterns and corresponding transcriptional profiles in human and canine osteosarcoma revealed strong correlations between key transcriptional patterns and DNA methylation measurements. For example, conserved green methylation clusters were positively correlated with cell cycle expression programs in both species while conserved orange methylation clusters were negatively correlated again in both species. These same methylation clusters displayed the opposite behavior when compared to the immune expression profiles highlighting the tradeoff between tumors with high proliferation and those with higher immune components also seen in Scott et. al. 2018. These findings have implication for developing potential biomarkers or other predictive measures for identifying tumors that might be more aggressive due to higher rates of proliferation or to track very low levels of immune cell infiltration that is often missed using standard pathology methods.

Next, we generated a new algorithm based on CIBERSORT and past use of Illumina 450K methylation profiles^16^ to predict immune cell abundances from whole genome bisulfite sequencing data. Our analysis shows a well-defined difference between immune cell profiles among human and canine osteosarcoma samples. Canine osteosarcoma show much higher presence of immune cells across the samples analyzed, average absolute abundance measures were higher for 6 out of 10 immune cells in the canine samples as well as for osteoclasts and MSC. Notably, we observed increased CD8+ T cell signature in canines (mean absolute abundance 0.44) that are relatively less abundant in human samples (mean absolute abundance 0.01). This observation is intriguing because even with higher immune cell infiltration in canine samples, this did not translate to increased survival in canines. Even though canine tumors are primarily driven by the proliferation signatures that are correlated with methylation patterns, we speculate that progression in canine tumors is predominantly driven by cell proliferation with immune cells potentially having less influence in the absence of immunomodulatory therapies.

We also observed high correlations between the global methylation patterns and immune cell abundance in both human and canine osteosarcoma. Strong correlations between single cell types, or cell types with correlated abundances, and methylation modules were not observed leading us to believe that global methylation patterns do not seem to be driven only by the abundance of immune cells in the microenvironment, but most likely reflect the proliferative and pro-survival status of the tumor cells. Inhibiting pro-survival pathways linked to osteosarcoma progression has therapeutic relevance. Conventional therapies—including the DNA-intercalating drugs doxorubicin and cisplatin^44^ and methotrexate, an anti-metabolite in combination with leucovorin—have very serious side effects, including decreased production of blood cells that leads to infection and damage to the bladder and kidney. Platinum-containing drugs often cause hearing loss. New drugs such as muramyl tripeptide^45^, rapamycin inhibitor^46^, and Trastuzumab^47^, show only marginal increases in overall survival, leaving survival rates still distressingly low. The dysregulated transcriptional programs that are coupled with methylation patterns can be restored by DNA- and chromatin-modifying drugs 5-Aza (5-Aza-2′-deoxycytidine (5-Aza or decitabine, a hypomethylating agent) and SAHA (Suberanilohydroxamic acid or vorinostat, a histone deacetylase inhibitor). Recently we showed that 5-AZA and SAHA treatment alter the transcriptional landscape of osteosarcoma cells towards one resembling RB expression^42^. Preclinical and early clinical studies combining 5-Aza with chemotherapies, peptide vaccines and immune checkpoint therapies found evidence that this treatment increases tumor suppressor expression and chemosensitivity. Numerous clinical studies have also reported that 5-aza improves the efficacy of antigen-directed immunotherapy in pediatric sarcomas[45]. Moreover, treating osteosarcoma cell lines with a combination 5-AZA and SAHA induced apoptosis, even in aggressive cell lines that are typically more resistant to treatment.

In summary, we have we have used genome-wide methylation patterns to reveal association between methylation clusters and conserved transcriptional programs of human and canine osteosarcoma. This first large-scale genome-wide DNA methylation study in both human and canine osteosarcoma revealed specific DNA methylation programs that are highly correlated to gene expression programs important to disease progression and survival in both human and canine osteosarcoma. Further, exploring the global DNA methylation patterns between different cell types we found that the stromal and immune cell types were in low abundance in human osteosarcomas while canine osteosarcoma samples showed relatively greater abundance of monocytes, T cells, and osteoclasts. These comparative studies on mechanisms that regulate conserved transcriptional programs in both human and canine osteosarcoma are critical to develop biomarkers^48^ and therapeutic targets.

## Methods

### Biospecimen collection and processing

Biospecimens were collected from newly diagnosed human patients or dogs with a confirmed diagnosis of appendicular osteosarcoma prior to treatment with cytotoxic chemotherapy drugs^8,11^.

Human specimens(n=24) were obtained from the University of Minnesota Biological Materials Procurement Network (UMN BioNet) under oversight of the University of Minnesota’s Institutional Review Board (IRB) with an Exemption-4 category, or from the Cooperative Human Tissue Network (CHTN), also with an IRB Exemption-4. Samples were de-identified and only a limited amount of patient information was provided. Sample collection was done using standardized protocols with a portion of the diagnostic biopsy, obtained as part of a medically necessary procedure flash frozen immediately in liquid nitrogen and stored at −86°C until they were assigned to the project.

Dog specimens (n=44) were obtained from dogs with naturally occurring primary appendicular tumors, recruited between 1999 and 2016. Samples were obtained from tissue biopsy or amputation surgeries that were part of standard treatment protocols and with owner consent under supervision by the appropriate Institutional Animal Care and Use Committees (University of Minnesota protocol numbers 0802A27363, 1101A94713, 1312-31131A) or the University of Colorado Institutional Review Board or Institutional Animal Care and Use Committee (AMC 635040202, AMC 200201jm, AMC 2002141jm, 02905603(01)1F, COMIRB 06-1008). Some of these canine tumor samples were flash frozen immediately in liquid nitrogen and stored at −86°C until they were assigned to the project; others were immediately placed in complete, sterile cell culture media consisting of Dulbecco’s Modified Eagle Media supplemented with 10% fetal bovine serum, 10 mM 4-(2-hydroxyethyl)-1-piperazineethanesulfonic acid (HEPES), and 100 µg/mL Primocin and transported to the lab by overnight courier at 4°C, and flash frozen at −86°C after examination and processing upon arrival. Transport did not have a meaningful effect on tumor viability based on gross examination and on the functional capability to establish viable, immortalized osteosarcoma cell lines from the samples.

Tumor tissues were removed from −86°C and sectioned to avoid areas of necrosis. Thirty mg of tissue were placed in 80 µL of phosphate buffered saline (PBS) solution and pulverized in a tissue homogenizer. Isolation of genomic DNA was done according to the manufacturer’s protocol using the QIAamp DNA Mini Kit from Qiagen.

### Bisulfite conversion, library preparation and target region capture

Illumina library preparation, bisulfited converted and bead capture was performed as specified in the Roche SeqCap Epi protocol outlined in Li et al^49^. Probes for the canine version of the SeqCap Epi were designed for canFam3.1 CpG islands as specified by the UCSC genome browser and CpG islands identified using EMBOSS cpgreport (http://www.bioinformatics.nl/cgi-bin/emboss/cpgreport) that were homologous to hg19. An average of 48 and 46 million 2×125 PE Illumina reads were generated for the human and canine samples.

### Bisulfite-seq data analysis

Illumina specific adapter sequences and low quality sequences were removed from raw sequencing data using Trimmomatic^50^. Reads were aligned to a bisulfite converted human (hg19) or canine (canfam3.1) genome using WALT^51^. Duplicate sequences were removed from alignment data using MethPipe^52^ duplicate-remover and bisulfite conversion rates were estimated for each sample using bsrate. All samples had conversion efficiencies > 99%. Sequencing depths for each sample were such that an average of 88.3% and 81.3% of targeted CpGs were sequenced at a 10X depth or higher. MethPipe methcounts was used to calculate methylation levels and read coverage at individual CpGs in each sample. A custom perl script was used to isolate CpGs with a read depth of 10 or greater and MethPipe roimethstats was used to summarize methylation levels in 500 bp windows, generated by BEDtools^53^ windowBed, along the capture targets in each genome. CpG methylation levels were expressed as a value between 0 and 1.

### Identification of homologous genomic regions and region to gene mapping

USCS utility liftOver with the appropriate genome wide alignment file was used to convert the canine genomic coordinates for each 500bp window from CanFam3.1 to hg19. and the human genomic coordinates for each 500bp window to GRCh38. Converting both sets of coordinates to GRCh38 resulted in the highest number of 500bp windows being conserved between the two species. Genomic regions were considered homologs if the converted GRCh38 coordinates were within 1,000bp of each other as calculated by BEDtools closest.

### Identification of highly correlated DNA methylation patterns with WGCNA, cluster membership and correlation analysis

WGCNA analysis was carried out via a custom R script. Methylation levels for homologous genomic regions were the starting input for WGCNA in each species. All homologous genomic regions with a variance greater than 0 were included in the WGCNA analysis. Soft thresholding power was calculated for each species separately using pickSoftThreshold (human power = 10, canine power = 14, sup figure). Clusters (aka modules) were generated using blockwiseModules with the following parameters for both species and the species-specific power value.

~~~
blockwiseModules(<inputMethmatrix>,
                  maxBlockSize = 1500,
                  power = 14|10,
                  TOMType = “signed”,
                  corType = “bicor”,
                  minModuleSize = 20,
                  mergeCutHeight = 0.25,
                  verbose = 3,
                  numericLabels = T,
                  saveTOMs=T,
                  saveTOMFileBase=<name>)
~~~

Summary methylation values for each cluster were obtained from the MEs element of the output from blockwiseModules. The correlation (KME) between the methylation level at each genomic region to every methylation cluster summary value was calculated using signedKME and a genomic region was determined to be a member of a specific cluster if the KME was >= 0.7. This classification allows individual genomic regions to be assigned to multiple clusters. Pearson correlations between summary methylation values for each cluster and other phenotypic measures (gene expression, age, survival time, immune cell abundances) were calculated using the WGCNA function cor and Student asymptotic p-values were calculated for each correlation value using corPvalueStudent and the number of samples for the multiple sample correction. Identical WGCNA analysis performed on a randomized human dataset resulted in the generation of 6 clusters that contained 113 DNA methylation regions and randomized canine data did not produce any clusters.

### Generation of custom signature file for CIBERSORT

BigWig files containing genome wide methylation levels from WGBS data for multiple replicates of 12 cell types were downloaded from the BLUEPRINT epigenome project. BigWig files were converted to bedGraph using the UCSC utility bigWigToBedGraph, sorted using BEDtools sort and mean methylation levels were calculated for the same 500bp windows used for the WGCNA analysis using BEDtools map -c 4 -o mean. These summarized methylation levels were used as the input for pairwise differential methylation tests between every cell type, 66 tests in all. Differential tests were performed using the limma^54^ package in R. The top 100 most differentially methylated regions, based on adjusted p-value, from all differential tests were isolated and the raw methylation values for each cell type for these regions were combined and used as input to CIBERSORT to generate the custom signature file. Raw methylation values for these same regions for all osteosarcoma samples were used as the mixture file input for CIBERSORT. Absolute Mode was used to quantify cell abundance to try and account for cell types that may be missing from our model but are in the tumor.

### Data Access and Sharing

All raw sequencing data, DNA methylation measurements, cluster membership and DNA methylation summary values have been submitted to GEO GSE149679

## Supporting information

BLUEPRINT_downloads

Canine methylation values 500bp regions

Canine pathway analysis

Canine cluster members

Canine clusters in each color

Average methylation for pure cell population for all diff meth regions

Human methylation values 500bp regions

Human pathway analysis

Human cluster members

Human clusters in each color

## Funding

Data analysis pipeline development was supported by Roche Sequencing Solutions, Madison. MB, PS, and TR are past or present employees of Roche Sequencing Solutions, Madison.

Work performed at University of Minnesota was supported in part by:

Zach Sobiech Fund for Osteosarcoma Research of the Children’s Cancer Research Fund;

National Cancer Institute/NIH grants R21 CA208529, R50 CA211249, and P30 CA077598;

Department of Defense (Congressionally Designated Medical Research Program) grant CA170218;

Morris Animal Foundation grant D13CA-032;

Karen Wyckoff Rein in Sarcoma Foundation;

Comparative Medicine Signature Program, College of Veterinary Medicine, University of Minnesota; Animal Cancer Care and Research Program, University of Minnesota.

## Acknowledgements

David Largaespada for his work with the Sobiech group and conversations about the project.

The authors acknowledge the Minnesota Supercomputing Institute (MSI) at the University of Minnesota for providing resources that contributed to the research results reported within this paper. URL: http://www.msi.umn.edu

## Author contributions

LM – analyzed the data, developed the story, wrote first draft of results, outlined intro and discussion.

MS – worked with Nimblegen to establish approach for methylation analysis and develop kits. Intimately involved with design and initial data analysis. Prepared all canine samples for the project.

PS – Nimblegen. Performed the work to develop and validate the methylation kits and completed QC of the data. Presented original poster that provided a template for analysis.

AC – Roche NimbleGen. Facilitated in the provision of sequence capture products, protocols and training requirements.

AD – UMGC. Involved in design of canine methylome kit. Provided oversight for methylation sequencing.

BA – UMGC. Involved in design of canine methylome kit. Performed technical aspects of bisulfite sequencing.

BC – prepared all human samples for the project.

KB – UMGC. Involved in design of canine methylome kit. Provided funding and oversight for the project.

LS – Sobiech group. Provided funding and helped with intellectual conceptualization of the project.

AS – Sobiech group. Provided funding and helped with intellectual conceptualization of the project.

SS – Sobiech group. Provided funding and helped with intellectual conceptualization of the project.

TR – Nimblegen. Designed canine methylome approach and kit (mapped, developed baits, etc.) Provided funding for the project.

JM – Sobiech group. Provided funding and helped with intellectual conceptualization of the project. Overall responsibility for coordinating and moving the project forward.

---

**Supplementary Figure 1:**
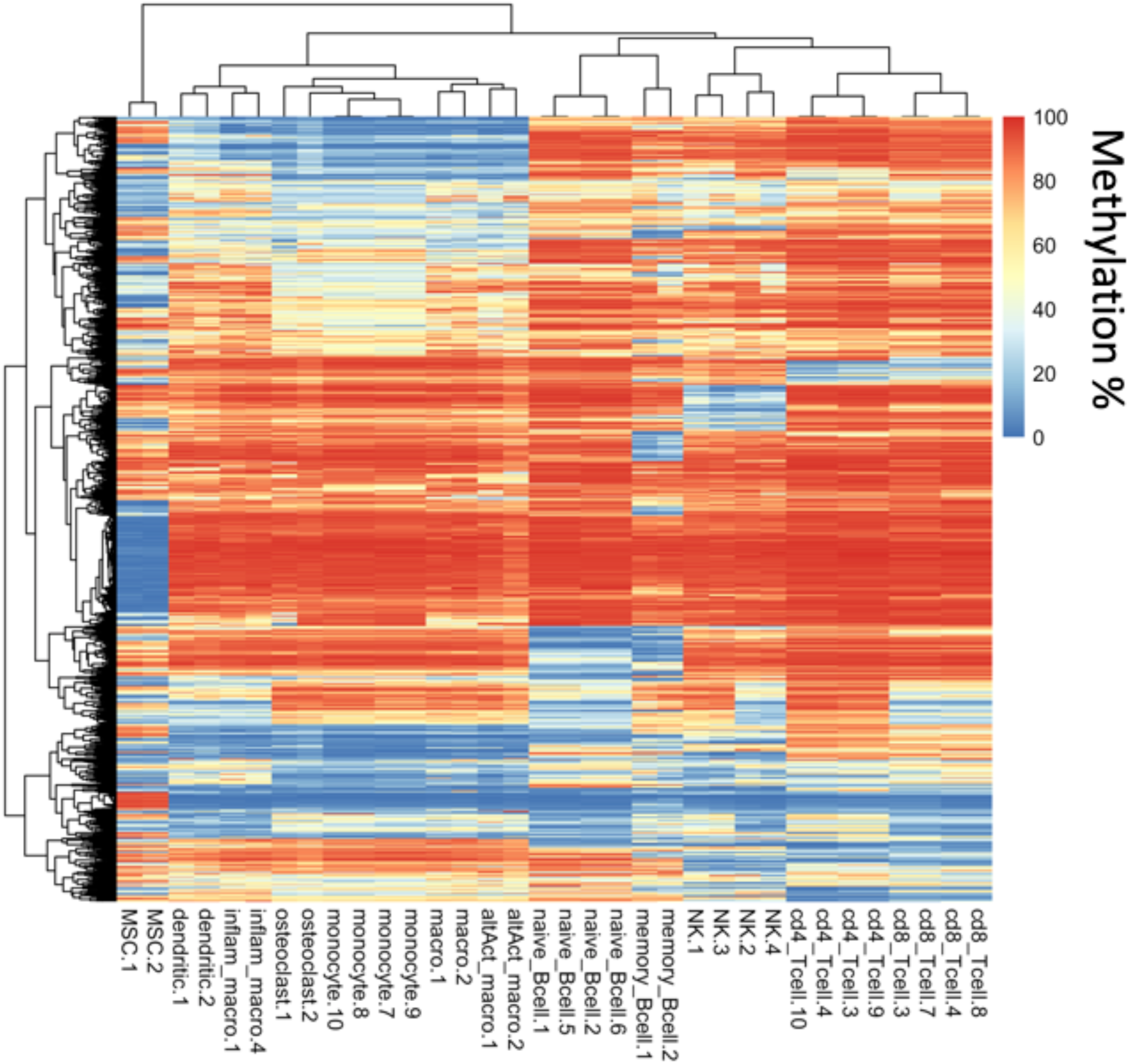
Differentially methylated regions from pure cell populations. Whole genome bisulfite sequencing data for 12 cell types of interest was obtained from BLUEPRINT. WGBS data was summarized to the same regions used to identify methylation modules and all pairwise differentially methylation tests were performed between all 12 cell types. The top 100 differently methylated regions in all pairwise comparisons were combined to generate the signature genes needed for CIBERSORT. Heatmaps give methylation values for each region for each pure cell sample for all100 top differentially methylated regions across all pairwise comparisons. High methylation values are red, low methylation values are blue.

**Supplementary Figure 2:**
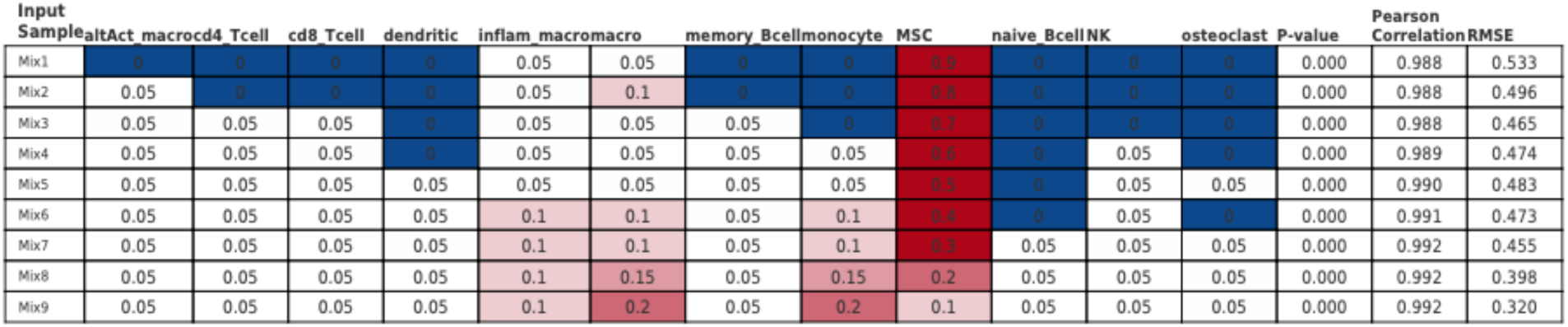
CIBERSORT is able to recreate known mixtures using custom signature genes. Whole genome bisulfite sequencing data for 12 cell types of interest was obtained from BLUEPRINT. WGBS data was summarized to the same regions used to identify methylation modules and all pairwise differentially methylation tests were performed between all 12 cell types. The top 100 differently methylated regions in all pairwise comparisons were combined to generate the signature genes needed for CIBERSORT. This signature gene file was able to 100% replicate the mixture of cell types that were generated computationally from the pure cell populations.

## Notes

### Competing Interest Statement

The authors have declared no competing interest.

### Summary of Updates

Spelling and grammatical fixes and some formatting changes. Fixed author names and affiliations.

